# Interaction forces reflect the perception of texture during active exploration

**DOI:** 10.64898/2026.06.17.732939

**Authors:** Neema Darabi, Benoit P. Delhaye, Sliman J Bensmaia, Stephanie E Palmer, Anton R Sobinov

## Abstract

We are constantly exploring the world around us through touch. Active touch depends on coordination of movement and force, yet the forces used during natural exploration and their relationship with perception is largely unexplored. Here we measured exploration forces together with fingertip motion while 17 participants explored 14 textures and rated their perceived hardness, slipperiness, or roughness. Exploration strategies differed systematically across tasks: hardness judgments involved relatively stationary pressing with larger and more variable normal forces, whereas slipperiness and roughness judgments relied more on sweeping movements. Within tasks, interaction forces covaried with perceptual ratings: harder textures elicited larger maximum tangential forces, consistent with diagonal pressing, while more slippery textures were explored with faster fingertip motion and lower forces. Estimated dynamic friction was negatively correlated with perceived slipperiness but not roughness. At the same time perceived roughness was strongly related to vibrations in the force, indicating distinct physical bases for these perceptual dimensions. These results show that humans actively tailor contact mechanics to perceptual goals during active exploration, supporting a sensorimotor account of texture perception.

**Significance statement:** Touch is usually studied as if the skin passively receives information, but in everyday life we actively move and press against objects to recognize them and learn what they feel like. This study measured both fingertip motion and contact forces while people freely explored textures and judged hardness, slipperiness, and roughness – the three of the most salient dimensions of tactile experience. The results show that people adjust how they move and press depending on what they want to perceive, and that different physical signals – friction and vibration – support different texture judgments. This work helps explain touch as an active sensorimotor process, with implications for neuroscience, haptics, robotics, and neuroprosthetics.

## Introduction

From stroking the hand of a loved one, to finding the power button on the back of a TV screen, we routinely use active touch to extract information about surfaces and objects. Human touch can discriminate remarkably fine surface differences down to a fraction of a millimeter, moreover, the way we move during haptic exploration depends on the property we are trying to judge (Skedung et al., 2013; Lieber and Bensmaia, 2019). Classic work on exploration strategies, termed exploratory procedures, showed that hand movements are not arbitrary: people select specific movements because they are especially effective for extracting specific object properties (Lederman and Klatzky, 1987). Tactile perception depends not only on the sensitivity of the skin, but also on how the hand inspects the world.

During texture exploration, contact forces between the skin and the surface provide the mechanical input that activates cutaneous mechanoreceptors and ultimately gives rise to tactile perception (Johnson et al., 2000; Birznieks et al., 2001; Bensmaïa and Hollins, 2003; Johansson and Flanagan, 2009; Weber et al., 2013; Jones and Smith, 2014; Lieber and Bensmaia, 2022). Work using active, passive, and pseudo-passive paradigms suggests that texture perception is affected by the mechanics and sensorimotor context of exploration, including movement (Hollins and Risner, 2000; Yoshioka et al., 2011; Pruszynski et al., 2018). Nevertheless, studies of active texture exploration have largely emphasized perceptual judgments, controlled passive stimulation, or exploratory kinematics, while the force patterns accompanying natural exploration remain poorly characterized (Smith et al., 2002; Callier et al., 2015). This gap matters because normal force, tangential force, friction, and force fluctuations all shape the mechanical signal delivered to the skin and can themselves relate to perceptual judgments. In particular, tangential force fluctuations have been linked to roughness judgments, and both normal and tangential components contribute to the perception of dynamic contact (Roberts et al., 2020; Gueorguiev et al., 2022). A fuller account of tactile exploration should therefore measure forces and movements together while subjects pursue distinct perceptual goals.

In this study, subjects explored textures while focusing on a single perceptual feature: slipperiness, roughness, or hardness, which are the most salient perceptual dimensions outside of temperature (Bensmaïa and Hollins, 2005; Callier et al., 2015). We measured normal and tangential interaction forces together with kinematics during exploration and recorded subjects’ perceptual ratings. We then asked whether characteristic force ranges emerge for each task, whether those forces vary systematically with the perceptual feature being judged, and which physical variables best predict perceptual reports. By linking interaction forces, movement, and perception during unconstrained exploration, this study aims to clarify how we actively sample surfaces to support different perceptual goals. This work further provides a quantitative framework for studying how interaction forces contribute to active texture perception.

## Results

Seventeen participants were tasked with rating how slippery, hard, or rough the presented texture is after exploring it with the pad of their index finger (Figure 1A). We have chosen fourteen textures that span a wide range of each of these perceptual axes: acrylic, bumpy polyester, denim, foam, grid upholstery, sueded cuddle, microsuede, rubber, 240 grit sandpaper, 800 grit sandpaper, snowflake fleece, tan upholstery, velvet, white foam. The textures were mounted on a large 30.5 by 30.5 cm plexiglass surface to encourage free and unconstrained exploration. They were also hidden from participants’ view by a cloth, so they were relying exclusively on somatosensory cues. Each texture was presented to each participant three times in each of the three perceptual blocks. To assess the rating consistency across participants, we compared the variability of the mean rating for each texture with a shuffled distribution in which texture identities were randomly mixed across all trials (Figure 1B). We have found that participants were consistent in their ratings, although slipperiness was the most variable dimension, consistent with previous findings (Hollins et al., 2000), with acrylic eliciting the least consistent ratings.

**Figure 1.**
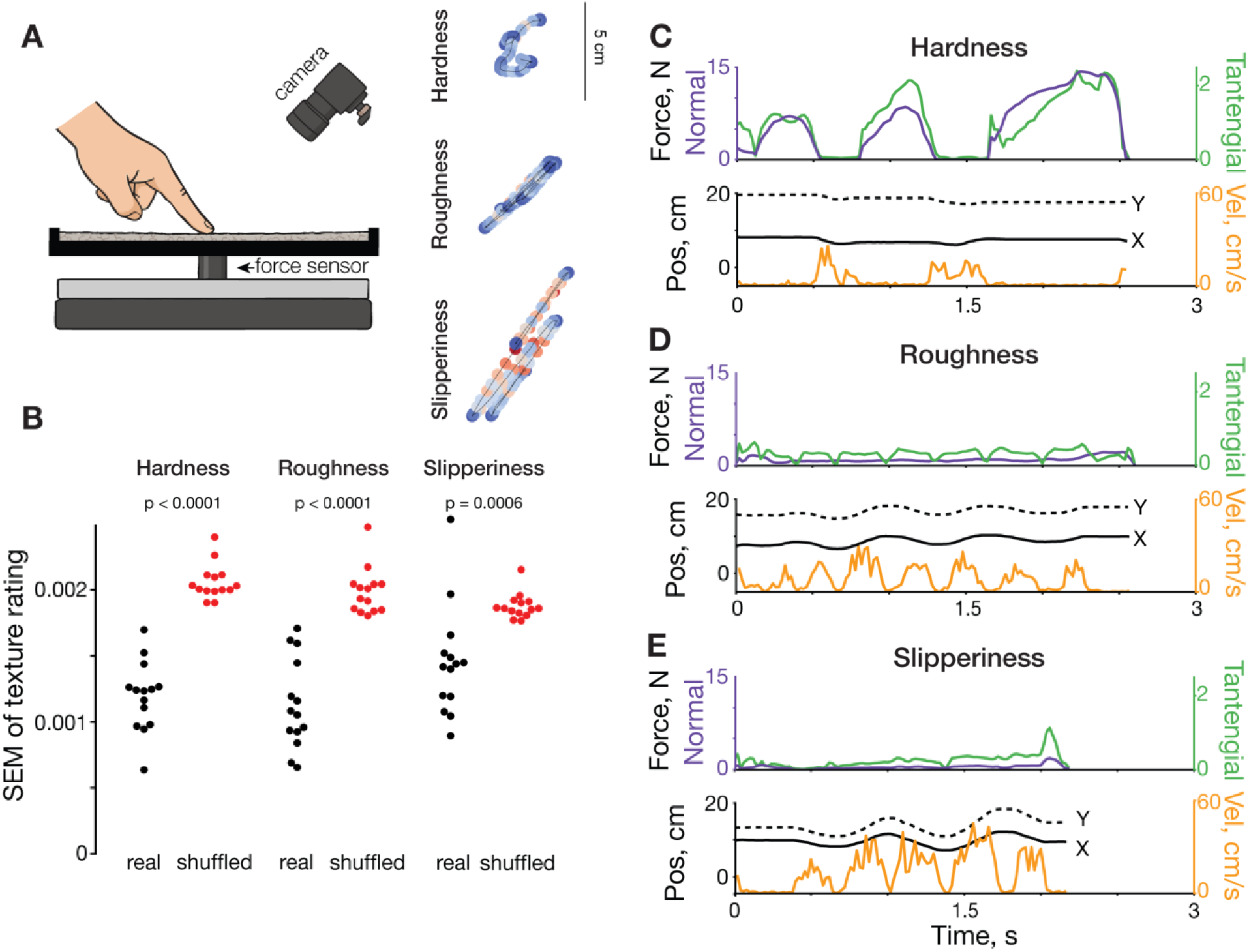
Experimental setup and data examples. **A**| Experimental setup and example fingertip position traces during a single trial (same trials as CDE). Color of each point corresponds to velocity. **B**| Rating consistency. The standard error of the mean of ratings within texture (‘real’) and when texture identity was shuffled between trials, pooled over all participants. Mann-Whitney U test, N=14 textures. **CDE**| Normal force, tangential force, position, and velocity traces during a single trial exploration of hardness,

In each trial, the textures were securely attached to a steel plate mounted on a force sensor, which allowed us to record the interaction forces as well as the movement of the finger using high-speed camera (Figure 1CDE). We have observed several stereotypical behavioral patterns during exploration of each perceptual feature. The exploration of hardness was often characterized by presses into the texture, resulting in simultaneously varying normal and tangential volitional force traces (Figure 1**Error! Reference source not found**.C). On those trials, fingertip position remained relatively constant, showing only slight velocity bursts when presses were initiated, or a finger was moved to a different spot. Slipperiness and roughness exploration involved swiping exploration strategy. Normal forces were characterized by the peaks in the beginning and end of each swipe, remaining largely constant between those (Figure 1DE). Tangential force had dips in its profile, when the sweeping finger changed the direction of movement, also reflected in periodic fingertip position and speed traces similar to previously reported ones (Callier et al., 2015). The exploration strategies appeared similar between roughness and slipperiness, and their ratings were negatively correlated (p=0.005, r=-0.70, Supplementary Figure 1).

Participants employed a large variety of forces during texture exploration (see Methods, Table 1). The normal forces, measured across all subjects and tasks, had a median of 0.79 N (IQR 0.29-2.10 N). The tangential forces were smaller, as expected, with a median at 0.49 N (IQR 0.20-1.09 N). The highest average normal forces of 1.7 N, 1.6 N, and 1.5 N were applied during exploration of hardness of white foam, bumpy polyester, and rubber, respectively. The peak forces estimated as 99^th^ percentile, however, were much larger and reached 34 N. The median tangential forces were similar between all textures, peaking around 5-14 N.

**Table 1.**
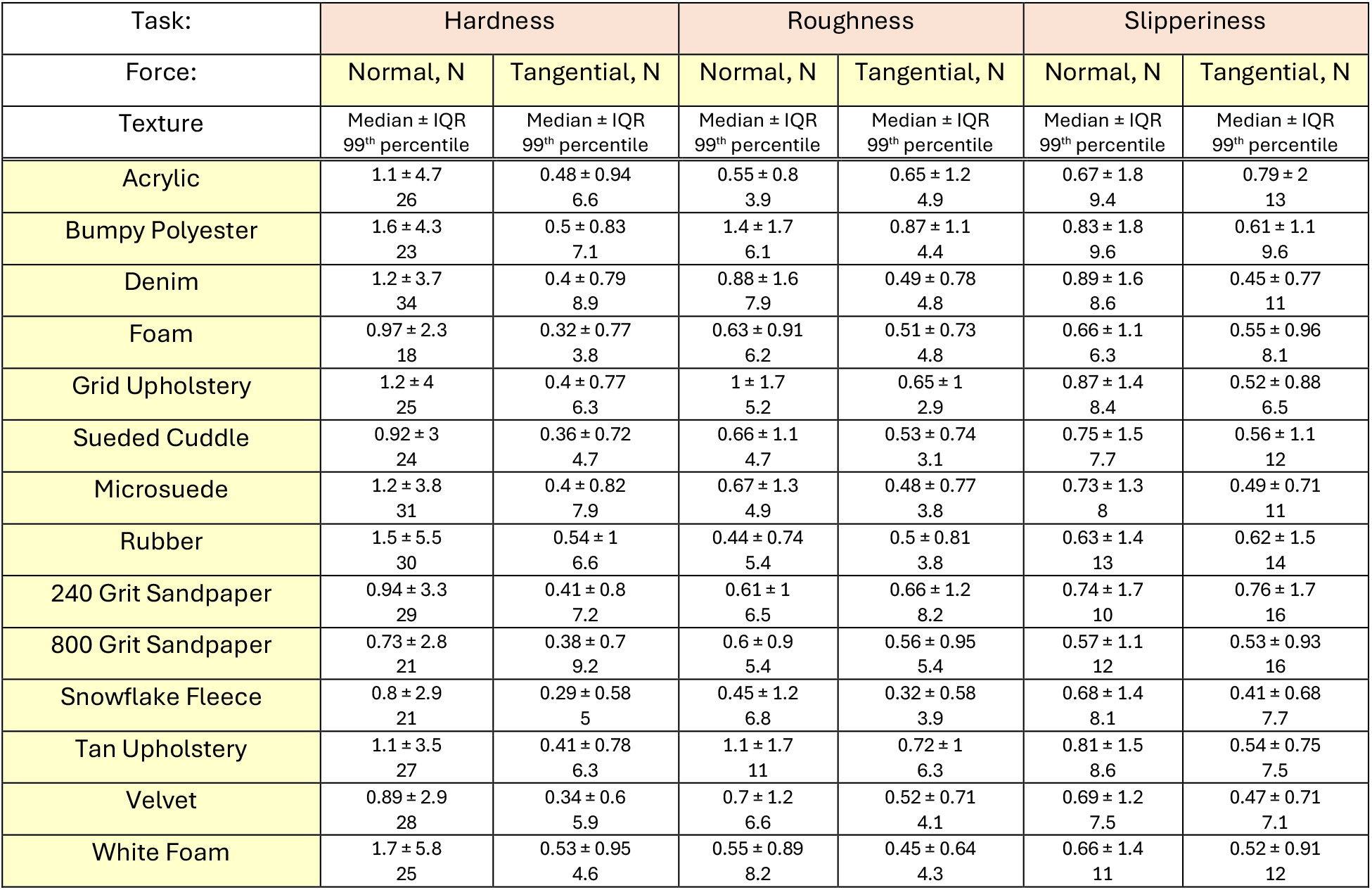
Mean forces used during texture exploration, organized by task, texture, and force dimension.

| Task: | Hardness |  | Roughness |  | Slipperiness |  |
| --- | --- | --- | --- | --- | --- | --- |
| Force: | Normal, N | Tangential, N | Normal, N | Tangential, N | Normal, N | Tangential, N |
| Texture | Median ± IQR<br>99<sup>th</sup> percentile | Median ± IQR<br>99<sup>th</sup> percentile | Median ± IQR<br>99<sup>th</sup> percentile | Median ± IQR<br>99<sup>th</sup> percentile | Median ± IQR<br>99<sup>th</sup> percentile | Median ± IQR<br>99<sup>th</sup> percentile |
| Acrylic | 1.1 ± 4.7<br>26 | 0.48 ± 0.94<br>6.6 | 0.55 ± 0.8<br>3.9 | 0.65 ± 1.2<br>4.9 | 0.67 ± 1.8<br>9.4 | 0.79 ± 2<br>13 |
| Bumpy Polyester | 1.6 ± 4.3<br>23 | 0.5 ± 0.83<br>7.1 | 1.4 ± 1.7<br>6.1 | 0.87 ± 1.1<br>4.4 | 0.83 ± 1.8<br>9.6 | 0.61 ± 1.1<br>9.6 |
| Denim | 1.2 ± 3.7<br>34 | 0.4 ± 0.79<br>8.9 | 0.88 ± 1.6<br>7.9 | 0.49 ± 0.78<br>4.8 | 0.89 ± 1.6<br>8.6 | 0.45 ± 0.77<br>11 |
| Foam | 0.97 ± 2.3<br>18 | 0.32 ± 0.77<br>3.8 | 0.63 ± 0.91<br>6.2 | 0.51 ± 0.73<br>4.8 | 0.66 ± 1.1<br>6.3 | 0.55 ± 0.96<br>8.1 |
| Grid Upholstery | 1.2 ± 4<br>25 | 0.4 ± 0.77<br>6.3 | 1 ± 1.7<br>5.2 | 0.65 ± 1<br>2.9 | 0.87 ± 1.4<br>8.4 | 0.52 ± 0.88<br>6.5 |
| Sueded Cuddle | 0.92 ± 3<br>24 | 0.36 ± 0.72<br>4.7 | 0.66 ± 1.1<br>4.7 | 0.53 ± 0.74<br>3.1 | 0.75 ± 1.5<br>7.7 | 0.56 ± 1.1<br>12 |
| Microsuede | 1.2 ± 3.8<br>31 | 0.4 ± 0.82<br>7.9 | 0.67 ± 1.3<br>4.9 | 0.48 ± 0.77<br>3.8 | 0.73 ± 1.3<br>8 | 0.49 ± 0.71<br>11 |
| Rubber | 1.5 ± 5.5<br>30 | 0.54 ± 1<br>6.6 | 0.44 ± 0.74<br>5.4 | 0.5 ± 0.81<br>3.8 | 0.63 ± 1.4<br>13 | 0.62 ± 1.5<br>14 |
| 240 Grit Sandpaper | 0.94 ± 3.3<br>29 | 0.41 ± 0.8<br>7.2 | 0.61 ± 1<br>6.5 | 0.66 ± 1.2<br>8.2 | 0.74 ± 1.7<br>10 | 0.76 ± 1.7<br>16 |
| 800 Grit Sandpaper | 0.73 ± 2.8<br>21 | 0.38 ± 0.7<br>9.2 | 0.6 ± 0.9<br>5.4 | 0.56 ± 0.95<br>5.4 | 0.57 ± 1.1<br>12 | 0.53 ± 0.93<br>16 |
| Snowflake Fleece | 0.8 ± 2.9<br>21 | 0.29 ± 0.58<br>5 | 0.45 ± 1.2<br>6.8 | 0.32 ± 0.58<br>3.9 | 0.68 ± 1.4<br>8.1 | 0.41 ± 0.68<br>7.7 |
| Tan Upholstery | 1.1 ± 3.5<br>27 | 0.41 ± 0.78<br>6.3 | 1.1 ± 1.7<br>11 | 0.72 ± 1<br>6.3 | 0.81 ± 1.5<br>8.6 | 0.54 ± 0.75<br>7.5 |
| Velvet | 0.89 ± 2.9<br>28 | 0.34 ± 0.6<br>5.9 | 0.7 ± 1.2<br>6.6 | 0.52 ± 0.71<br>4.1 | 0.69 ± 1.2<br>7.5 | 0.47 ± 0.71<br>7.1 |
| White Foam | 1.7 ± 5.8<br>25 | 0.53 ± 0.95<br>4.6 | 0.55 ± 0.89<br>8.2 | 0.45 ± 0.64<br>4.3 | 0.66 ± 1.4<br>11 | 0.52 ± 0.91<br>12 |

Different participants used different ranges of forces, both normal and tangential (Supplementary Figure 2). The median normal force in the four participants with the highest forces was 4.94 times bigger than those of the participants with the lightest touch, in the hardness task (Supplementary Figure 2A, ANOVA p<0.001).

Similar relationship existed in the slipperiness task, and for tangential forces. To avoid inter-subject differences in applied forces influencing the comparison between texture forces and their ratings, we z-scored forces using the mean force used by each subject in subsequent analyses.

### Force differences during perceptual tasks

To determine whether participants adjusted their exploratory forces according to the perceptual feature being evaluated, we compared z-score-adjusted forces across tasks (Figure 2). Participants applied larger median normal forces when judging hardness than when judging slipperiness or roughness (Figure 2A). They also showed greater variability in normal force during hardness judgments (Figure 2C).

**Figure 2.**
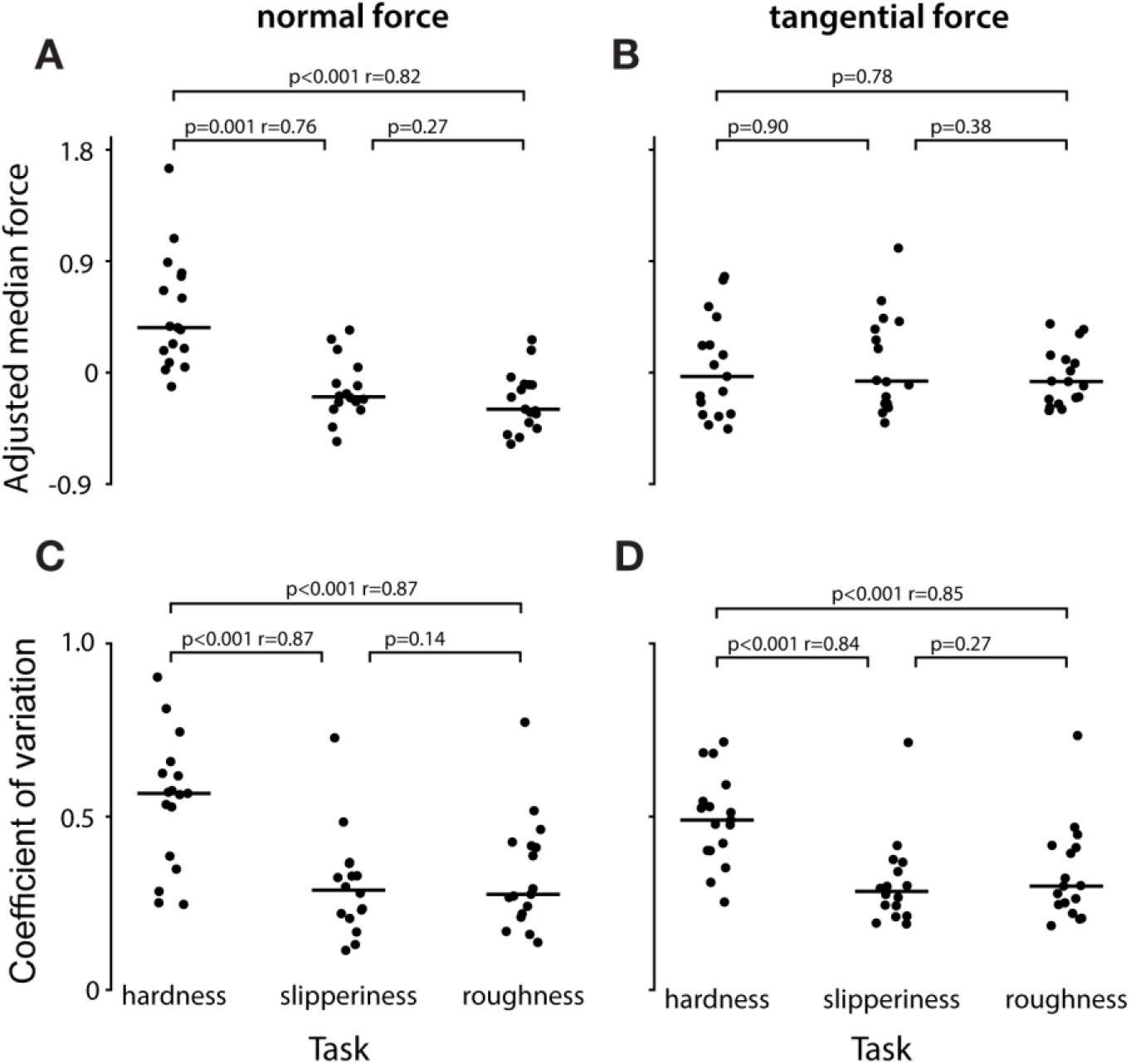
Effect of task on the applied forces. **A**| Adjusted median normalforce per subject during hardness, slipperiness, and roughness trials. **B**| Same as A, but for tangential force. **CD**| Same as AB but coefficient of variation. Median force is normalized by z-scoring each subject’s median force with the subject’s average force across all tasks. P-values represent Wilcoxon signed-rank test significance, and r-values – rank-biserial correlation. N=17 participants.

In the introductory example, we showed that participants pressed into the texture to judge its hardness (Figure 1B). Here, we found that this pressing was often diagonal, producing tangential forces comparable to those used during slipperiness trials (Figure 2B). In contrast, when judging slipperiness or roughness, participants used a sweeping strategy and kept tangential force relatively constant, as indicated by its lower variability than during hardness trials (Figure 2D). In summary, hardness exploration was characterized by strong diagonal pressing, whereas slipperiness and roughness exploration involved more consistent tangential force.

### Force correlates with texture ratings

Having shown that participants adapted their exploration strategies to the task, we next asked whether their movements and forces also reflected texture perception. To address this question, we used regression to predict ratings from eight kinematic and kinetic variables describing texture exploration: median, maximum, and absolute rate of change of total tangential force, median, maximum, and absolute rate of change of normal force, and mean and max fingertip speed (See Methods, Table 2). Fingertip speed was, as expected, a poor predictor of hardness ratings (Figure 3A). Instead, the strongest predictor was maximum tangential force: participants pushed harder textures away from themselves more forcefully than softer textures (Figure 3C). By contrast, neither maximum nor median normal force significantly predicted hardness rating (Figure 3B and Methods). Together, these results suggest that tangential force plays a central role in hardness perception, consistent with the idea that humans assess hardness by pressing into textures diagonally.

**Table 2.** Predictors of perceptual judgments in each task. The table lists the predictors, the cross-validated pseudo-R^2^ of the model, and one-sided permutation test p-value assessing how many shuffled distributions out of 1000 shuffles outperformed the model. Color highlights the type of the regressor: tangential force in green; normal force in red; kinematics in purple. Significant relationships (p<0.05) were highlighted in bold.

| Regressor | Hardness |  | Slipperiness |  | Roughness |  |
| --- | --- | --- | --- | --- | --- | --- |
| | $R^2$ | p-value | $R^2$ | p-value | $R^2$ | p-value |
| Index Fingertip Speed, Mean | -0.77 | 0.98 | **0.91** | **0.0001** | **0.44** | **0.0001** |
| Index Fingertip Speed, Max | -0.11 | 0.15 | **0.73** | **0.0001** | **0.44** | **0.0001** |
| Normal Force, Median | -0.73 | 0.96 | -0.16 | 0.19 | **0.24** | **0.012** |
| Normal Force, Max | -0.82 | 0.98 | **0.58** | **0.0001** | **0.26** | **0.01** |
| Normal Force Rate, Median | -0.31 | 0.58 | -0.37 | 0.73 | -0.23 | 0.32 |
| Tan. Force, Median | **0.155** | **0.022** | **0.37** | **0.002** | 0.007 | 0.068 |
| Tan. Force, Max | **0.39** | **0.002** | **0.66** | **0.0001** | -0.11 | 0.15 |
| Tan. Force Rate, Median | -0.35 | 0.66 | **0.21** | **0.008** | -0.12 | 0.16 |

**Figure 3.**
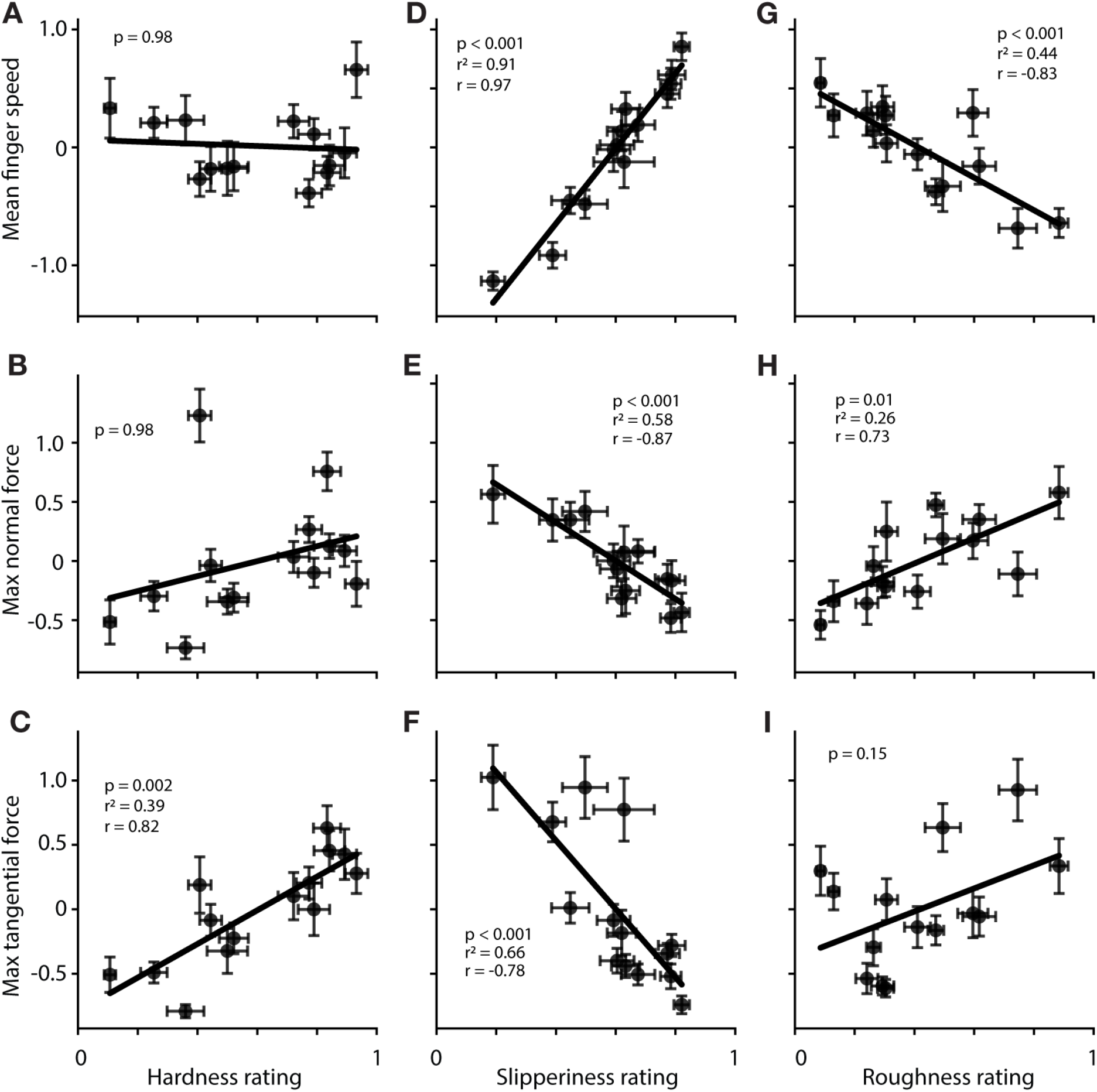
Modulation of kinematics and interaction forces with perception rating. **A**| Median adjusted fingertip speed during texture exploration vs hardness rating. **B**| Same as A for peak normal force. **C**| Same as A for peak tangential force. **DEF**| Same as ABC for slipperiness rating. **GHI**| Same as ABC for roughness rating. Whiskers indicate standard error of the mean computed on inter-subject variability; p-value was computed against shuffled distribution; r^2^– cross-validate pseudo-r^2^; r – in-sample correlation; N=14 textures.

Fingertip velocity was an extremely strong correlate of slipperiness rating (Figure 3D). Participants also exerted lower tangential forces when exploring more slippery textures (Figure 3F), consistent with previous studies showing that tangential force varies with texture friction (Smith and Scott, 1996). In addition, we found that normal force also decreased with increasing slipperiness (Figure 3E). This coordinated reduction in both force components may reflect an exploratory strategy that preserves sensitivity to frictional differences across textures. Because the coefficient of friction is defined as tangential force divided by normal force, joint modulation of these forces may help maintain informative friction-related signals. Specifically, variability in force production generally increases with force magnitude, therefore, reducing interaction forces may improve the reliability of slipperiness estimates for slippery textures. Overall, participants explored more slippery surfaces with faster movements and lower interaction forces, likely to maintain perceptual sensitivity.

The relationships between roughness ratings and behavioral variables largely mirrored those observed for slipperiness, although the effects were weaker (Figure 3GHI). Participants explored rougher textures with slower fingertip movements (Figure 3G) and greater normal forces (Figure 3H). Although tangential force was not a significant predictor of roughness ratings, it showed a trend in the expected direction (Figure 3I). Together, these results suggest that roughness and slipperiness judgments rely on partly overlapping exploratory strategies, with participants adjusting movement speed and interaction forces according to surface properties.

### Force vibrations are related to texture ratings

Moving the digit across the texture produces vibrations in it and the skin, activating distinct mechanoreceptors. To determine whether these vibrations were related to the perception of texture, we calculated the power of 5 frequency bands: 5-25 Hz, 25-50 Hz, 50-100 Hz, 100-400 Hz, and 400-1000 Hz – in trials at least 2 seconds long. We have selected these bins to roughly separate the ranges for different mechanoreceptors in glabrous skin (Johansson and Flanagan, 2009). Then, we computed Spearman correlations between that power and the perceptual rating of the texture, for each participant, and pooled across all participants (Figure 4). Intermediate-frequency vibrations were strongly predictive of hardness ratings, whereas neither low-frequency (<25 Hz) nor high-frequency (>400 Hz) vibrations showed comparable relationships (Figure 4A).

**Figure 4.**
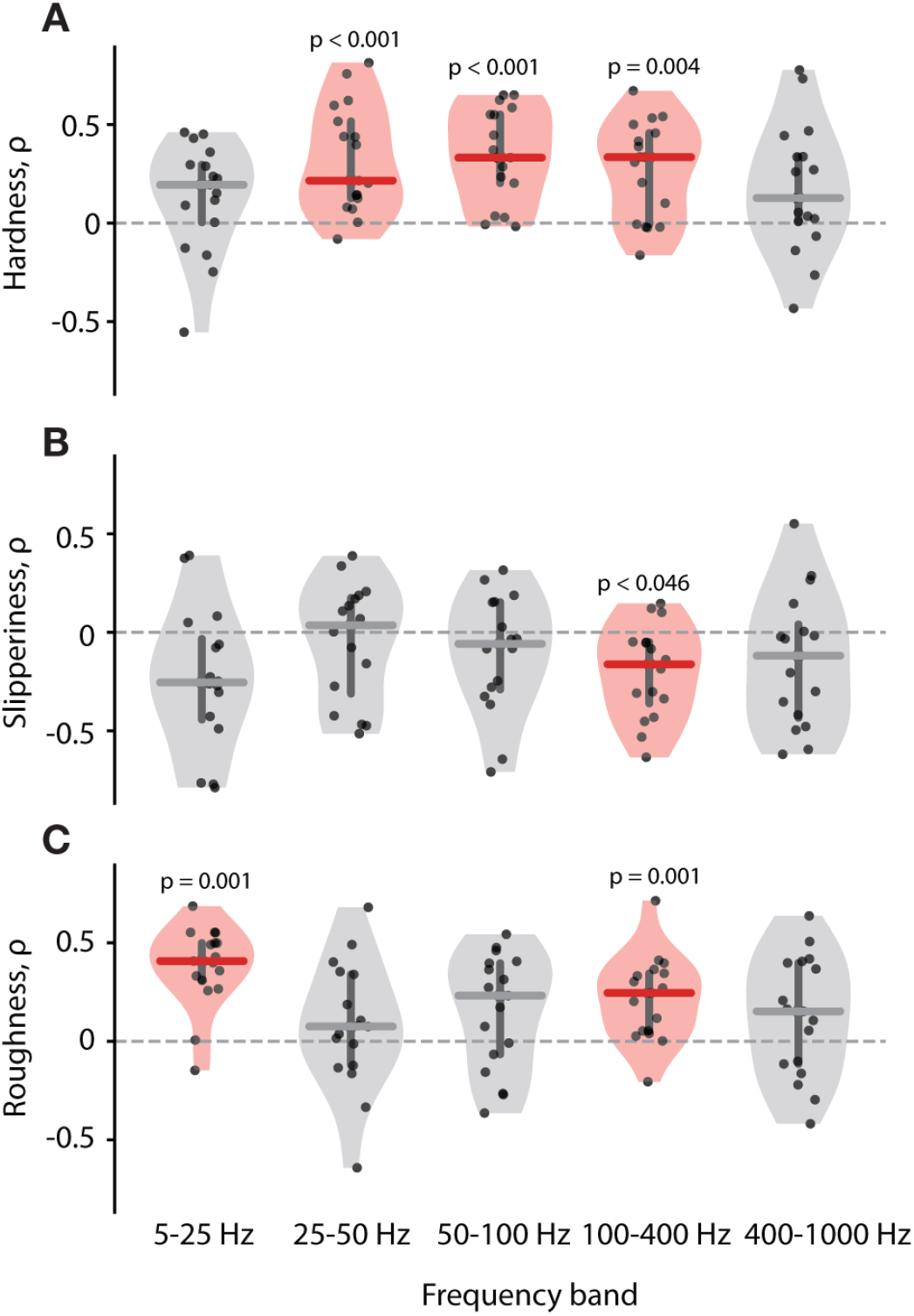
Relationships between power of force vibration at different bands and perceptual rating. **A**| Hardness rating. Each dot corresponds to a correlation computed in one subject (see Supplementary Materials). Two-tailed Wilcoxon signed-rank P-values were calculated on pooled data (N=230), then corrected using Benjamini-Hochberg for 5 frequency bands. **B**| Same as A for slipperiness. N=223. **C**| Same as A for roughness. N=238. Horizontal lines correspond to median, vertical ones – to first and third quartile. Significant relationships (p<0.05 after correction) highlighted in red.

Surprisingly, slipperiness was only weakly related to vibratory power, with significance limited to the 100-400 Hz band (Figure 4B). Roughness, on the other hand, was strongly related to low (5-25 Hz) frequency vibrations and, to a lesser extent, with vibrations in the 100- 400 Hz range (Figure 4C). Interestingly, the correlations in 100-400 Hz range had negative correlation with slipperiness and opposite – positive one – with roughness. Together, these results suggest that different perceptual dimensions of texture are supported by distinct spectral features of contact-induced vibrations during free exploration.

### Dynamic friction predicts perceived slipperiness

We then asked whether perceived slipperiness and roughness were related to an intrinsic physical property of the texture: its friction. The dynamic friction is calculated by dividing the tangential force by the normal one, while the digit is moving. In each trial, we identified periods of fingertip motion in line with the distribution of swiping speeds reported previously (Callier et al., 2015). For those periods, we computed the median ratio of tangential to normal force as an estimate of the coefficient of friction between the finger and the texture. Estimated values ranged from 0.3 to 1.8, broadly matching those reported in the literature (Smith and Scott, 1996; Savescu et al., 2008; Vilhena et al., 2023). Slipperiness ratings were strongly negatively correlated with texture friction (Figure 5A), showing that perceived slipperiness is closely tied to friction during unconstrained exploration. In contrast, roughness ratings did not show the same relationship with friction (Figure 5B), highlighting roughness as a distinct perceptual dimension linked to a different physical property of contact.

**Figure 5.**
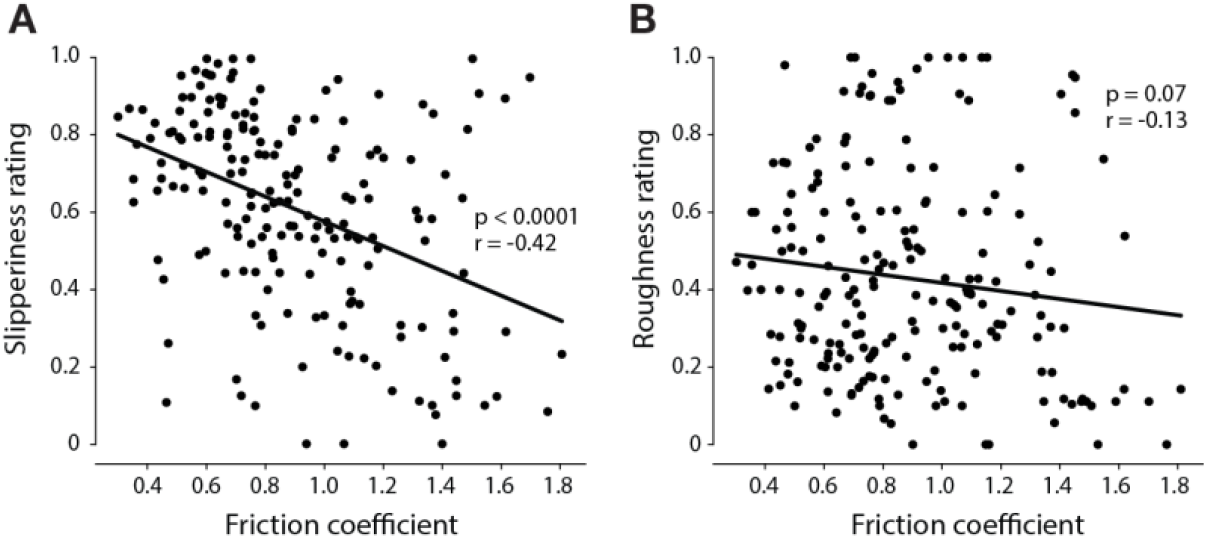
Relationships between perceptual rating and friction of a texture. **A**| Normalized slipperiness rating of a texture vs its friction coefficient. N=190 trials. **B**| Normalized roughness rating of a texture vs its friction coefficient. N=204 trials.

## Discussion

Our results show that active texture exploration is systematically shaped by perceptual goals. Participants did not use a single generic strategy across tasks. Instead, judgments of hardness were associated with larger and more variable interaction forces, whereas judgments of slipperiness and roughness relied more on sweeping movements of the fingertip. At the same time, participants differed markedly in the absolute forces they used, yet the task structure became clear after within-subject normalization. This combination of strong inter-subject variability and consistent within-subject modulation suggests that active touch is governed not by a fixed motor template, but by individualized exploratory policies that are nevertheless organized around common perceptual goals. This interpretation fits well with previous work showing that haptic exploration depends on the property being judged, as well as with more recent observations that texture-directed finger movements vary both with task and with the surface being explored (Lederman and Klatzky, 1987; Callier et al., 2015). However, here we show it during active unconstrained exploration.

The clearest task-dependent contrast was between hardness and slipperiness exploration. Hardness exploration was characterized by stronger loading, greater normal-force variability, and relatively stationary fingertip position, consistent with a pressing-based strategy for probing compliance. Importantly, the variable that best tracked perceived hardness was not normal force, but tangential force: participants pushed harder textures away from themselves more strongly than softer ones. That interpretation is consistent with prior work showing that exploratory movements in softness discrimination are tuned to stimulus compliance and that people spontaneously adjust contact force to optimize performance (Kaim and Drewing, 2011; Dhong et al., 2019). More broadly, tactile signals during object interaction are known to reflect the mechanics of skin deformation, including both indentation and lateral strain, which makes it plausible that hardness judgments depend on the combined loading pattern rather than on normal force alone (Hollins and Risner, 2000; Weber et al., 2013; Lieber and Bensmaia, 2022). Hardness ratings were also related to vibrations across a broad range of frequencies, suggesting that perceived hardness reflects the frequency-dependent dynamic stiffness of the material (Higashi et al., 2019).

Slipperiness judgments depended on a different exploratory strategy. Here, fingertip velocity was the strongest behavioral correlate of perceptual rating, and both normal and tangential forces decreased as textures were perceived as more slippery. The simultaneous decrease in both forces is likely related to an attempt at maintaining sensitivity to the relative change in the forces, which define the dynamic friction of the texture. We show that the slipperiness rating is closely related to the friction of the texture, which agrees well with the previous experiments performed in more restricted environments (Johansson and Westling, 1984; Smith and Scott, 1996).

At the same time, roughness did not exhibit the same relationship with friction as slipperiness, despite similarities in exploratory kinematics and the anticorrelation between their ratings. Instead, roughness was more strongly related to force vibrations, particularly in the low-frequency and 100– 400 Hz bands. Low-frequency vibrations may arise in part from the exploratory strategy itself, as the digit moves across coarse surface features and generates slow fluctuations in contact force (Shao et al., 2016). By contrast, the 100-400 Hz band overlaps the range of greatest vibrotactile sensitivity and will engage rapidly adapting tactile afferents (Makous et al., 1995). The relationship between roughness and vibrations in this band is therefore consistent with prior work showing that texture-induced vibrations, particularly those represented through Pacinian-weighted signals, contribute to perceived roughness (Bensmaïa and Hollins, 2003). Together, these findings suggest that roughness and slipperiness, although behaviorally related, depend on partially distinct mechanical cues: friction for slipperiness and spectrally specific force vibrations for roughness (Hollins and Risner, 2000). Thus, roughness and slipperiness appear to rely on overlapping exploratory behaviors but distinct mechanical signals, with roughness more strongly tied to vibration and slipperiness more closely tied to friction (Wiertlewski et al., 2011; van Kuilenburg et al., 2015; Willemet et al., 2021).

Although participants differed substantially in absolute force levels, the preservation of task-dependent structure after normalization suggests that these differences reflect individualized exploratory preferences rather than noise variability. At the same time, our measurements were limited to force, whereas pressure may also be relevant for texture perception, and could have been more equal between the participants (Westling and Johansson, 1984; Lederman and Klatzky, 1993; Derler et al., 2013). However, estimating pressure would require tracking the evolving contact patch throughout exploration because finger pad contact area changes dynamically with load and contact conditions (Dzidek et al., 2017). Combining behavioral measurements with high-resolution tactile arrays or contact imaging may therefore help determine whether distributed pressure, in addition to gross force, contributes to perceptual judgments in future research.

## Methods

### Participants

The methodology used in this experiment was approved by the University of Chicago Institutional Review Board for Human Use (IRB23-1405). 17 participants, all undergraduate or graduate students at the University of Chicago (age 20-29, 7F, 1 left-handed), performed the experiment. No participant reported any tactile deficits or any sensorimotor neurological deficits.

### Experimental protocol

In this study, participants were told to explore a texture and report a single number describing how rough, hard, or sticky the texture feels. Subjects were seated behind a black curtain to hide their hand and the texture. Before each trial, subjects were told to develop a rating of one of the three tactile features. Ratings could take on any range of numbers but needed to be proportional so that a texture three times as rough got a rating three times as large. When instructed to begin, subjects extended their arm outwards to explore the texture with their dominant hand’s index fingertip. Subjects explored textures until they had a rating in mind. They were allowed to use any set of strategies to develop this rating. After exploration of each texture was completed, the subject rated the texture feature. All ratings were normalized by the maximum rating in each perceptual task for each subject. After reporting the rating, they returned their arm to the armrest, and the texture was swapped for a new one, thus concluding the trial.

The fourteen textures were presented in random order three times, per textural feature. Each of the three feature tasks contained forty-two trials. All subjects completed each task except one who only completed two, having to leave due to unforeseen circumstances outside the experiment.

### Data collection

Textures were attached to 30.5 × 30.5 × 0.3 cm Plexiglas panels, allowing ample space for natural kinematic exploration strategies. To measure the forces used by subjects during exploration, the plexiglass panels were fixed to a square piece of metal with two screws. The metal support was mounted on top of an ATI Mini45 Titanium Six-Axis Force/Torque Sensor, placed on top of another metal layer. A square layer of rubber was also placed below the metal at the bottom to dampen external noise. The force sensor measured analog signals which were converted to normal and tangential forces in Newtons, using a force sensor-specific calibration matrix provided by the manufacturer (Calibration: SI-60-3; Method: WI-FTP-026, DAQ Calibration Instructions). We measured the total force placed on the texture, regardless of position, at 2000 Hz.

To track finger movement in two-dimensional space, a high-speed Blackfly S (BFS-U3-16S2C- CS) was placed about 50 cm above the texture, recording at 80 frames per second. A neural network from the open-source toolbox MMPose (https://github.com/open-mmlab/mmpose) tracked the wrist, the metacarpophalangeal (MCP), proximal interphalangeal (PIP), and distal interphalangeal (DIP) joints of the index finger, the thumb MCP joint, and the index fingertip in each frame of recording. Custom code converted pixels to mm and accounted for the camera angle. Global time markers indicating the beginning of force data and image data were used to synchronize the image tracking and force measurements.

### Data processing

We analyzed the three linear dimensions of volitional force, one being normal to the textures and the other two being tangential. Each force dimension was individually processed using a median filter (20 ms kernel size) and a low-pass Butterworth filter (order 2). To normalize the signal, we first identified periods of no contact by comparing the standard deviation of force in the first and last 100 data points (.05 s). The mean force of the segment with lower standard deviation was subtracted from the entire trace and multiplied the result by -1 so that baseline forces were centered at 0 N and contact resulted in positive force values. Then, negative forces, which reflect the spring-like behavior of the force sensor, were removed. The rest of the analysis included only the period between the first and the last instance of non-zero force. We identified this period by scanning the full force trace and defined points of contact as those where normal force exceeded 20 times the lower baseline standard deviation described above. The data was binned using the median of 20-ms windows. Although force recordings were set to 2000 Hz, the true sampling rate varied throughout trials. 0.22% of data points had a gap greater than 0.02 seconds with the subsequent recording, resulting in 2.56% of all 20-ms bins missing data and not being used in the analysis. These points were interpolated when needed for visualization.

The two components of force tangential to the texture were combined into one tangential force vector, resulting in one normal and one tangential component of volitional force in our analysis. The normal and tangential forces used by all subjects are summarized by task and texture in Table 1.

### Kinematic analysis

To analyze the kinematic properties of exploration, an MMPose hand pose and keypoint detection algorithm was run on each frame (pose estimator config: rtmpose-m_8xb256-210e_hand5-256×256; keypoint detector config: rtmdet_nano_320-8xb32_hand). This tracked the wrist, the MCP, PIP, and DIP joints of the index finger, the thumb MCP joint, and the index fingertip of any hand detected in the camera frame. We only tracked keypoints during the period of exploration, defined above. To this end, we reidentified the beginning and end of exploration for each trial, determined from force data. To synchronize force and image data, we calculated a time offset using global sync events producing an image and force timestamp for each trial. For the first 8 subjects, image timestamps were corrupted due to a frame buffer issue. We fixed this by modeling the timing error using data from subjects 9-17, generating a piecewise linear function via basin hopping minimization algorithm. The resulting image timing estimates introduced minimal error (mean residual of 5.45 * 10^-7^ s ± 0.087 s), acceptable given the 80-fps camera recording.

After determining the exploration period in terms of image frames, we determined the keypoints which belonged to the exploring hand, in instances where the neural network detected multiple hands. We manually labeled a bounding box using the four corners of the explored texture and in each frame, we only kept data from the hand with index fingertip inside the bounding box. Verified manually, this method accurately determined the exploring hand. The exploring hand’s keypoint data, for the period of exploration, was then filtered by keypoint score, followed by median filtering (5 frame, or about 62.5 ms kernel width) and Gaussian filtering (with a standard deviation of 0.5 frames), using interpolation in frames with low keypoint scores. The filtered data was then binned using 20 ms windows.

### Kinetic analysis

To compare the ratings between the subjects, we have normalized them to the respective ranges of ratings within a subject. Similarly, we have adjusted the measured forces by subtracting the mean and dividing by standard deviation.

To identify the behavioral features describing the perceptual judgements, we have selected 8 features describing individual trials: median, maximum, and absolute rate of change of tangential force, median, maximum, and absolute rate of change normal force, and mean and maximum index fingertip speed. We then used cross-validated regression to predict the rating of each texture using each one of these variable (Table 2).

### Vibration analysis

For each trial, contact was detected from the normal force after subtracting the pre-contact baseline, using a threshold defined as 10 times the standard deviation of the baseline force. Trials with contact durations shorter than 2 s were excluded. For valid trials, the final 2 s of the exploration period were extracted, mean-centered, and transformed into the frequency domain using Welch’s power spectral density estimate. Band-limited RMS amplitudes were then computed by integrating spectral power within five predefined frequency bands: 5-25, 25-50, 50-100, 100-400, and 400-1000 Hz. For each subject, trials were averaged by texture, and Spearman correlations were computed between each frequency-band amplitude and the corresponding perceptual ratings, separately for hardness, slipperiness, and roughness. The resulting subject-level correlation coefficients were then pooled across subjects for visualization and inference. In the pooled analysis, the distribution of subject-level Spearman correlations was plotted for each condition and frequency band, and significance was assessed using a two-tailed Wilcoxon signed-rank test against zero, with Benjamini-Hochberg correction across frequency bands within each condition.

## Acknowledgements

The authors would like to thank the participants for their time. This work was supported by the National Institute of Neurological Disorders and Stroke, R35 NS122333.

## Data availability

The data analyzed during this study are going to be made available on an online server upon publication https://uchicago.box.com/s/mgpe4qj8xq875hg4m14j8liz22mdo5ls.

## Code availability

All analysis and plotting scripts are going to be made available in a public repository https://github.com/SobinovLab/texture_force_paper

## Supplementary Materials

**Supplementary Figure 1.**
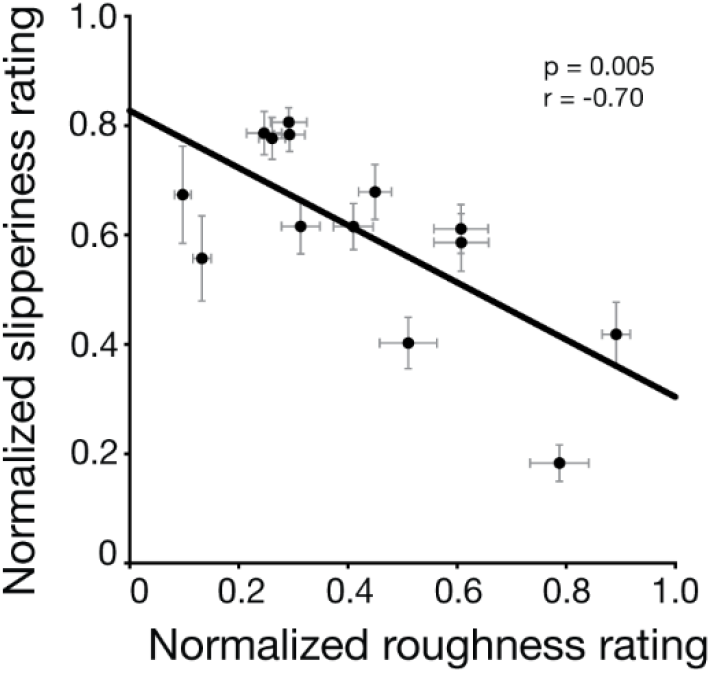
Relationship between slipperiness and roughness normalized rating for each texture.

**Supplementary Figure 2.**
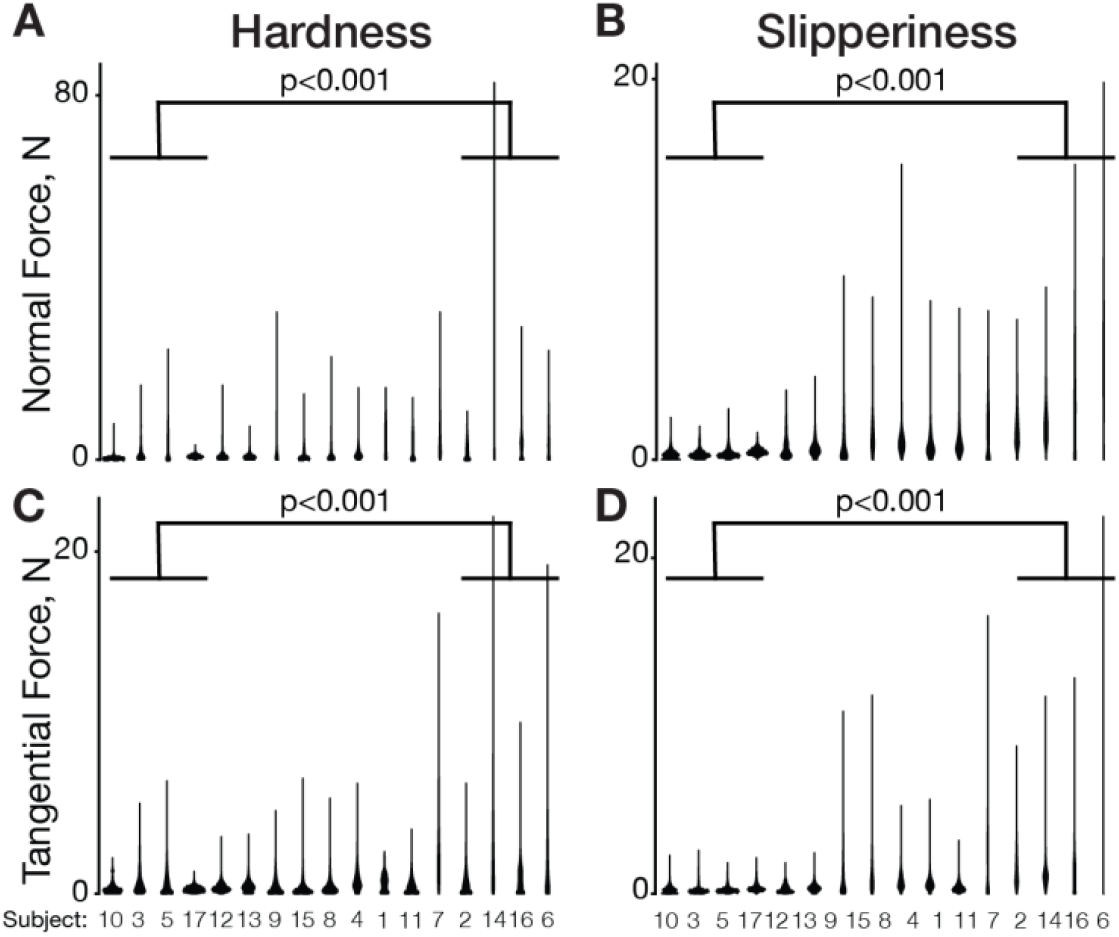
Distribution of normal and tangential forces for each task and subject. Columns correspond to hardness and slipperiness, while rows – to normal and tangential force. The subjects were sorted in the order of increasing mean normal force. Significance according to ANOVA test is indicated between the 4 subjects who used the highest and lowest overall forces.

## References

Bensmaïa S, Hollins M (2005) Pacinian representations of fine surface texture. Percept Psychophys 67:842–854.

Bensmaïa SJ, Hollins M (2003) The vibrations of texture. Somatosens Mot Res 20:33–43.

Birznieks I, Jenmalm P, Goodwin AW, Johansson RS (2001) Encoding of Direction of Fingertip Forces by Human Tactile Afferents. J Neurosci 21:8222–8237.

Callier T, Saal HP, Davis-Berg EC, Bensmaia SJ (2015) Kinematics of unconstrained tactile texture exploration. J Neurophysiol 113:3013–3020.

Derler S, Süess J, Rao A, Rotaru G-M (2013) Influence of variations in the pressure distribution on the friction of the finger pad. Tribol Int 63:14–20.

Dhong C, Miller R, Root NB, Gupta S, Kayser LV, Carpenter CW, Loh KJ, Ramachandran VS, Lipomi DJ (2019) Role of indentation depth and contact area on human perception of softness for haptic interfaces. Sci Adv 5:eaaw8845.

Dzidek BM, Adams MJ, Andrews JW, Zhang Z, Johnson SA (2017) Contact mechanics of the human finger pad under compressive loads. J R Soc Interface 14:20160935.

Gueorguiev D, Lambert J, Thonnard J-L, Kuchenbecker KJ (2022) Normal and tangential forces combine to convey contact pressure during dynamic tactile stimulation. Sci Rep 12:8215.

Higashi K, Okamoto S, Yamada Y, Nagano H, Konyo M (2019) Hardness Perception Based on Dynamic Stiffness in Tapping. Front Psychol 9 Available at: https://www.frontiersin.org/journals/psychology/articles/10.3389/fpsyg.2018.02654/full [Accessed June 4, 2026].

Hollins M, Bensmaïa S, Karlof K, Young F (2000) Individual differences in perceptual space for tactile textures: Evidence from multidimensional scaling. Percept Psychophys 62:1534–1544.

Hollins M, Risner SR (2000) Evidence for the duplex theory of tactile texture perception. Percept Psychophys 62:695–705.

Johansson RS, Flanagan JR (2009) Coding and use of tactile signals from the fingertips in object manipulation tasks. Nat Rev Neurosci 10:345–359.

Johansson RS, Westling G (1984) Roles of glabrous skin receptors and sensorimotor memory in automatic control of precision grip when lifting rougher or more slippery objects. Exp Brain Res 56:550–564.

Johnson KO, Yoshioka T, Vega-Bermudez F (2000) Tactile functions of mechanoreceptive afferents innervating the hand. J Clin Neurophysiol Off Publ Am Electroencephalogr Soc 17:539–558.

Jones LA, Smith AM (2014) Tactile sensory system: encoding from the periphery to the cortex. WIREs Syst Biol Med 6:279–287.

Kaim L, Drewing K (2011) Exploratory Strategies in Haptic Softness Discrimination Are Tuned to Achieve High Levels of Task Performance. IEEE Trans Haptics 4:242–252.

Lederman SJ, Klatzky RL (1987) Hand movements: A window into haptic object recognition. Cognit Psychol 19:342–368.

Lederman SJ, Klatzky RL (1993) Extracting object properties through haptic exploration. Acta Psychol (Amst) 84:29–40.

Lieber JD, Bensmaia SJ (2019) High-dimensional representation of texture in somatosensory cortex of primates. Proc Natl Acad Sci 116:3268–3277.

Lieber JD, Bensmaia SJ (2022) The neural basis of tactile texture perception. Curr Opin Neurobiol 76:102621.

Makous J, Friedman R, Vierck C (1995) A critical band filter in touch. J Neurosci 15:2808–2818.

Pruszynski JA, Flanagan JR, Johansson RS (2018) Fast and accurate edge orientation processing during object manipulation Verstynen T, ed. eLife 7:e31200.

Roberts RD, Loomes AR, Allen HA, Di Luca M, Wing AM (2020) Contact forces in roughness discrimination. Sci Rep 10:5108.

Savescu AV, Latash ML, Zatsiorsky VM (2008) A technique to determine friction at the finger tips. J Appl Biomech 24:43–50.

Shao Y, Hayward V, Visell Y (2016) Spatial patterns of cutaneous vibration during whole-hand haptic interactions. Proc Natl Acad Sci 113:4188–4193.

Skedung L, Arvidsson M, Chung JY, Stafford CM, Berglund B, Rutland MW (2013) Feeling Small: Exploring the Tactile Perception Limits. Sci Rep 3:2617.

Smith AM, Chapman CE, Deslandes M, Langlais J-S, Thibodeau M-P (2002) Role of friction and tangential force variation in the subjective scaling of tactile roughness. Exp Brain Res 144:211–223.

Smith AM, Scott SH (1996) Subjective scaling of smooth surface friction. J Neurophysiol 75:1957– 1962.

van Kuilenburg J, Masen MA, van der Heide E (2015) A review of fingerpad contact mechanics and friction and how this affects tactile perception. Proc Inst Mech Eng Part J J Eng Tribol 229:243– 258.

Vilhena L, Afonso L, Ramalho A (2023) Skin Friction: Mechanical and Tribological Characterization of Different Papers Used in Everyday Life. Materials 16:5724.

Weber AI, Saal HP, Lieber JD, Cheng J-W, Manfredi LR, Dammann JF, Bensmaia SJ (2013) Spatial and temporal codes mediate the tactile perception of natural textures. Proc Natl Acad Sci U S A 110:17107–17112.

Westling G, Johansson RS (1984) Factors influencing the force control during precision grip. Exp Brain Res 53:277–284.

Wiertlewski M, Lozada J, Hayward V (2011) The Spatial Spectrum of Tangential Skin Displacement Can Encode Tactual Texture. IEEE Trans Robot 27:461–472.

Willemet L, Kanzari K, Monnoyer J, Birznieks I, Wiertlewski M (2021) Initial contact shapes the perception of friction. Proc Natl Acad Sci U S A 118:e2109109118.

Yoshioka T, Craig JC, Beck GC, Hsiao SS (2011) Perceptual Constancy of Texture Roughness in the Tactile System. J Neurosci 31:17603–17611.

